# Embedding of exogenous B cell epitopes on the surface of UreB structure generates a broadly reactive antibody response against *Helicobacter pylori*

**DOI:** 10.1101/2021.02.09.430551

**Authors:** Junfei Ma, Shuying Wang, Qianyu Ji, Jingxuan Qiu, Qing Liu

## Abstract

Since *Helicobacter pylori* (*H. pylori*) resistance to antibiotic regimens is increased, vaccination is becoming an increasingly important alternative therapy to control *H. pylori* infection. UreB, FlaA, AlpB, SabA, and HpaA proteins of *H. pylori* were previously proved to be used as candidate vaccine antigens. Here, we developed an engineered antigen based on a recombinant chimeric protein containing a structural scaffold from UreB and B cell epitopes from FlaA, AlpB, SabA, and HpaA. The multi-epitope chimeric antigen, named MECU, could generate a broadly reactive antibody response including antigen-specific antibodies and neutralizing antibodies against *H. pylori* urease and adhesins. Moreover, therapeutic immunization with MECU could reduce *H. pylori* colonization in the stomach and protect the stomach in BALB/c mice. This study not only provides a promising immunotherapy to control *H. pylori* infection, but also offers a reference for antigen engineering against other pathogens.

## 1. Introduction

*Helicobacter pylori* (*H. pylori*) infection has been shown to be associated with a series of gastric diseases including chronic gastritis, peptic ulcers, and gastric malignancies^1, 2^. The major antibiotic-based triple regimen has been proven to cause increased antimicrobial resistance^3^, which results in a reduced eradication rate. Therefore, vaccination could be a more effective and safer immunotherapy to control *H. pylori* infection.

The key functional proteins of bacteria, which play an important role in invading and colonizing the host, are generally selected as vaccine antigens. In *H. pylori*, urease could neutralize the stomach acidity and promote chemotaxis by decomposing urea, which is conducive to the bacterial survival^4, 5^. Urease subunit beta (UreB) was widely used as a vaccine antigen against *H. pylori* infection, whose B cell epitopes and CD4^+^ T cell epitopes of UreB had been identified in the previous studies^6–11^. Besides, some other proteins of *H. pylori* were also developed as vaccine antigens. A linear B cell epitope of the movement-related protein flagellin A (FlaA) was also identified by experiment^12^. *H. pylori* adhesins including sialic acid-binding adhesin (SabA), hop family adhesin AlpB (AlpB), and *H. pylori* adhesin A (HpaA) are known to be promising candidate vaccine antigens against *H. pylori*^13^. Their possible conservative adhesion domains have also been reported in the previous studies^14–16^. The immune response induced by above antigens could inhibit the bacterial chemotaxis and adhesion.

Since the immune response induced by a single protein antigen is limited, the emphasis on vaccine development has moved to the generation of recombinant multi-epitope vaccines. A multi-epitope vaccine consists of B and T epitopes from several different antigens in a reasonable order with flexible linkers^17^. Compared to the single protein antigen vaccine, the multi-epitope vaccine could activate broadly reactive antibodies, which could lead to an effective immunoreaction blocking multiple pathogenic channels for the pathogen control^18^. A variety of multi-epitope vaccines against *H. pylori* could induce high levels of specific antibodies against multiple antigens which are the sources of epitopes^19–21^. The construction of multi-epitope vaccine antigen is undoubtedly the creation and synthesis of a new protein. The rationality of epitope assembly determines the expression certainty, stability, and degradation of the constructed multi-epitope antigen. Proper presentation of antigens, which can efficiently induce the immune system, is strongly dependent on the optimal structural stability of the vaccine construct^22^. Although computer-aided design could engineer immunogens according to the available structural information, most studies remained at design stage^23, 24^, suggesting that the string-of-beads structural rationality of multi-epitope vaccine antigen designed by computer aid needs to be proven.

In terms of identifying appropriate engineered antigens to obtain optimal vaccine response, considerable work has focused on structural vaccinology, in which immunogens are rationally engineered using available structural information^22, 25^. The methods have been developed to transplant epitopes to scaffold proteins for structural stabilization. The engineered immunogens could present one or more key epitopes or immunogenic domains to induce epitope-specific antibodies or generate a broadly reactive antibody response. In recent studies, Roundleaf bat HBV core antigen (RBHBcAg), calcium binding antigen 43 homolog (Cah) and *H. pylori* ferritin were respectively selected as the scaffold proteins for displaying key epitopes or immunogenic domains of pathogens including hepatitis B virus, Shiga toxin-producing *Escherichia coli* and *Borrelia burgdorferi*^26–28^. The structure-based engineered immunogen has higher structural stability and more reasonable epitope exposure compared to epitope-tandem immunogens, which means that it could generate a broader and more persistent antibody response.

In this paper, an engineered immunogen named MECU was designed for displaying multiple B cell epitopes from FlaA, SabA, AlpB and HpaA on the structural surface of UreB, a scaffold protein which was widely used as *H. pylori* antigen. MECU promotes the production of potent antigen-specific antibodies and neutralizing antibodies against *H. pylori* urease and adhesins. The therapeutic vaccination of MECU showed that it could generate a broadly antibody response and reduce bacterial loads in stomach for controlling *H. pylori* infection in BALB/c mice. The construction of MECU could be a candidate vaccine against *H. pylori* infection and provide a reference for immunogen engineering against other pathogens.

## 2. Materials and Methods

### 2.1 Bacteria and animals

The mouse-adapted *H. pylori* strain SS1 was from our lab collection. *H. pylori* was cultured on Columbia agar plates enriched with 7% new-born calf serum, polymyxin B (5 μg/mL), trimethoprim (5 μg/mL), and vancomycin(10 μg/mL) under microaerophilic conditions (5% O_2_, 10% CO_2_, and 85% N_2_) at 37 °C for 3-5 days.

Specific pathogen-free (SPF) BALB/c mice, 5–6 weeks of age, 14±2 g, were purchased from Jiesijie (Shanghai, China). The mice were allowed 1 week to adapt to the environment before starting the experiments. This study was approved by China Ethics Committee.

### 2.2 Construction and expression of the multi-epitope chimeric antigen

The B cell epitopes in *H. pylori* adhesins (HpaA, AlpB, and SabA) were predicted by BepiPred 2.0 Server^29^ and a B cell epitope in FlaA were obtained from the Immune Epitope Database (IEDB). The selected epitopes were listed in Table 1. Among them, three epitopes from adhesins contain the previously reported adhesion domains^14–16^. Based on the structure of UreB (PDB ID: 1e9zB), five positions (UreB_4-14_, UreB_38-64_, UreB_463-477_, UreB_516-534_, UreB_552-569_) in the non-core surface regions at the N-terminal and C-terminal were selected for epitope replacements. After replaced by five B cell epitopes with the available linkers “KK”, “GS”, “GGS”, and “GGGS”, UreB was converted into multi-epitope chimeric UreB (MECU).

**Table 1.**
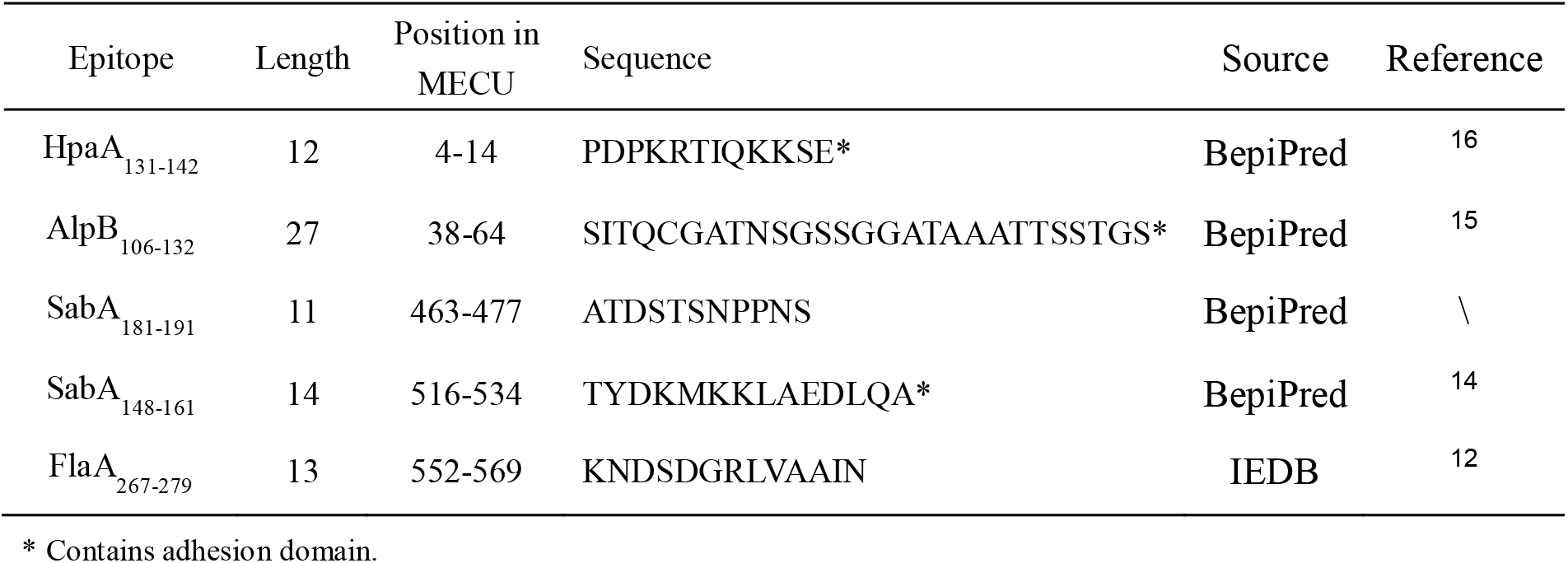
The selected B cell epitopes from HpaA, AlpB, SabA, and FlaA.

The amino acid sequence of the constructed MECU was submitted to the I-TASSER Server^30^ for structure prediction. ProSA-web, RAMPAGE, and Verify 3D sever were used for evaluating the quality of the predicted MECU structure. The Z-score calculated by ProSA-web could assess the overall quality of the predicted structure^31^. The main Ramachandran plot from RAMPAGE was used for calculating phi–psi torsion angles for each amino acid in the vaccine structure^32^. The Verify 3D Server could score the compatibility of the predicted structure model with the amino acid sequence^33^. Besides, the structural alignment of MECU and UreB was performed by TM-align tool^34^.

The amino acid sequence of MECU was submitted to the Jcat tool for Codon optimization, which could achieve maximum expression in *Escherichia coli* (*E. coli*) system^35^. The optimized DNA sequence was synthesized and cloned into the pET30a plasmid by Sangon Biotech(Shanghai, China). Finally, the recombinant plasmid pET30a-MECU was transformed into *E. coli* BL21(DE3) to express the MECU protein. The MECU protein was purified by affinity chromatography (Ni-IDA-Sefinose Column, Sangon Biotech, China). After that, the purified MECU protein was separated on 12% SDS-PAGE and further probed with mouse polyclonal anti-His (Sigma, USA) and rabbit polyclonal anti-*H. pylori* (GeneTex, USA) antibodies by western blotting with purified UreB as the reference.

### 2.3 Subcutaneous immunization with MECU

SPF BALB/c mice were randomly divided into 3 groups (n=5), and were vaccinated subcutaneously 3 times at 7-day intervals with 100 μg of the purified MECU, UreB or BSA in complete Freund’s adjuvant (FA, Sigma, USA) on day 0 and in incomplete FA on days 7 and 14. The pure proteins MECU, UreB or BSA were used as immunogens in the last booster immunization on days 21. Serum was collected at 7 days after the last vaccination to determine the specific antibodies.

### 2.4 Determination of specific antibodies after subcutaneous immunization

ELISA plates were coated with 1 μg/well of each antigen (UreB, FlaA, SabA, AlpB or HpaA) respectively at 4 ◻ overnight. After washing, the plates were blocked with 5% (m/V) skim milk for 2 h. The diluted serum samples (1:1000) were added to the antigen-coated plates and incubated for 1 h. After washing, a proper dilution of HRP-conjugated goat anti-mouse IgG (Sigma, USA) was added to the plate and incubated for 1 h. Finally, tetramethylbenzidine (TMB) was added and incubated at room temperature for 15 min. The reaction was then stopped with 2 M H_2_SO_4_. The absorbance was measured at 450 nm using a microplate reader.

### 2.5 Detection of neutralizing antibodies against *H. pylori* urease and adhesins

Serum IgG antibodies were purified by protein G column chromatography (GE Healthcare, USA). The purified IgG antibodies were detected by SDS-PAGE. To determine neutralizing antibodies against *H. pylori* urease, *H. pylori* urease (2μg in 50 μL, Creative Enzymes, USA) was incubated with purified IgG antibodies (64 μg/well) in 96-well microtiter plates overnight at 4 °C. After that, 100 μL of 50 mM PBS containing 500 mM urea, 0.02% phenol red, and 0.1 mM dithiothreitol (DTT) was added to each well. The absorbance was measured at 550 nm. Percentage inhibition of urease activity = [(activity without antibodies − activity with antibodies)/(activity without antibodies)] × 100 %.

To determine the neutralizing antibodies against *H. pylori* adhesins, the AGS human gastric cancer cell adhesion assay was carried out on *H. pylori* SS1 using a CFU counting method. Approximate 2×10^5^ AGS cells were seeded in 12-well plate per well overnight. Approximate 2×10^7^ CFU *H. pylori* cultures were incubated with 40 μL of purified IgG antibodies (100 μg/mL) and slightly shaken at room temperature for 2 h. AGS cells were washed three times with PBS, and the *H. pylori* incubates were added to the wells at a multiplicity of infection of 100. The mixtures were incubated in a CO_2_ incubator at 37 °C for 2 h. After incubation, the wells were washed 3 times with PBS containing 1% saponin. After the mechanical treatment, the mixtures were plated onto Columbia agar for bacteria counting. Percentage inhibition of adhesion = [(CFU without antibodies − CFU with antibodies)/(CFU without antibodies)] × 100 %.

### 2.6 Construction of CTB-MECU for oral immunization

To enhance the immunogenicity of oral MECU vaccine, cholera toxin subunit B (CTB), a widely used mucosal adjuvant, was added to the N-terminus of MECU and UreB with a flexible linker “DPRVPSS”. After amplification, cloning and transformation, CTB-MECU and CTB-UreB were also expressed by *E. coli* BL21 (DE3). The proteins were purified by affinity chromatography and assessed by SDS-PAGE. Their immunoreactivity was evaluated by western blotting.

CTB could bind intestinal epithelial cells and antigen presenting cells through monosialotetrahexosylganglioside (GM1) receptors, which then mediates antigen entry into the cell^36^. GM1-ELISA was used to demonstrate the adjuvanticity of CTB component in CTB-MECU as previously described^37^. Briefly, ELISA plates were coated with GM1 ganglioside at 4 ◻ for 12 h. After washing, ELISA plates were locked by incubating with 5% (m/V) skim milk for 2 h. The CTB-MECU, CTB-UreB, CTB, MECU, UreB or BSA proteins were added to the plates and incubated at 37 ◻ for 2 h. After washing, a proper dilution (1:1000) of anti-CTB mouse monoclonal antibody (Sigma, USA) was added to the plates and incubated at 37 ◻ for 1 h. After washing, HRP-conjugated goat anti-mouse IgG (Invitrogen, USA) was added to the plates ang incubated at 37 ◻ for 1 h. Substrate tetramethylbenzidine was then added and incubated for 15 min. The absorbance was measured at 450 nm.

### 2.7 Infection and therapeutic vaccination

SPF BALB/c mice (male, 5-6 weeks old) were infected with *H. pylori* SS1(10^9^ CFU/mouse) intragastrically, four times within the span of two weeks. The *H. pylori*-infected mice were randomly divided into 5 groups (n=5). Considering the different immunization methods, two groups were vaccinated intraperitoneally 3 times at 7-day intervals with 100 μg of the purified MECU or UreB in FA for four times at 1-week intervals. Other two groups were vaccinated intragastrically with 100 μg of antigen (CTB-MECU or CTB-UreB) in 0.2 M sodium hydrogen carbonate buffer (200 μL) for four times at 1-week interval. The last group was vaccinated with both FA in PBS intraperitoneally and CTB intragastrically as a control. Two weeks after the final immunization, the mice were sacrificed and examined. The whole therapeutic vaccination procedure was showed in Fig. 6A.

### 2.8 Assay of specific IgG in serum and SIgA in stomach mucosa

ELISA plates were coated with 5μg/mL *H. pylori* lysates at 4 ◻ overnight. To determine specific IgG, the antisera were collected and diluted 1:1,000 in PBS. To determine secretory IgA (SIgA), one-fourth of stomach tissue was homogenized in 1 mL of PBS containing 0.1 mM Phenylmethanesulfonyl fluoride (PMSF). The supernatant was collected and diluted 1:5 in PBS. An HRP-conjugated goat anti-mouse IgG or an HRP-conjugated goat anti-mouse IgA (Sigma, USA) was used as the secondary antibody.

### 2.9 Examination of *H. pylori* colonization in stomachs

To examine the *H. pylori* colonization in stomachs, one-fourth of stomach tissue was weighed and homogenized in 1 mL of PBS. Serial 10-fold dilutions of the stomach homogenate were plated on Columbia agar supplemented with 7 % new-born calf serum and *H. pylori* selective supplement (Oxoid, UK). After cultured for 4-6 days at 37 ◻, colonies were counted and the number of CFU per stomach was calculated.

### 2.10 Gastric histology

One fragment of stomach tissue was fixed with formalin, embedded in paraffin and stained with hematoxylin and eosin (HE) according to the standard procedure^38^.

### 2.11 Cytokine production

To determine cytokine production, the splenic lymphocytes were isolated and cultured (2×10^5^ cells/well) with *H. pylori* lysates (5 μg/mL) in 12-well plates at 37 ◻ for 72 h. The culture supernatants were collected for the determination of IFN-γ, IL-4, and IL-17 using ELISA kits (Jiang Lai Biotech, Shanghai, China) according to the manufacturer’s instructions.

### 2.12 Statistical analyses

All independent experiments carried out in this study and showed in figures were biological replicates. All data were analyzed with GraphPad Prism software using One-way ANOVA. P<0.05 was considered as statistically significant (* p < 0.05, ** p < 0.01, *** p < 0.001; ns, not significant).

## 3. Results

### 3.1 Construction and expression of MECU antigen

Based on the structural framework of UreB, five B cell epitopes from four antigens associated with adhesion or motion(HpaA, AlpB, SabA, and FlaA) were chosen to construct the MECU antigen (Fig. 1A). On the one hand, the epitopes located in the non-core regions next to the N-terminal and C-terminal of MECU primary structure. On the other hand, the insertion of exogenous epitopes did not affect the integrity of UreB own epitopes (Fig. 1B). This made the antigenicity of UreB fully embodied in MECU.

**Fig. 1.**
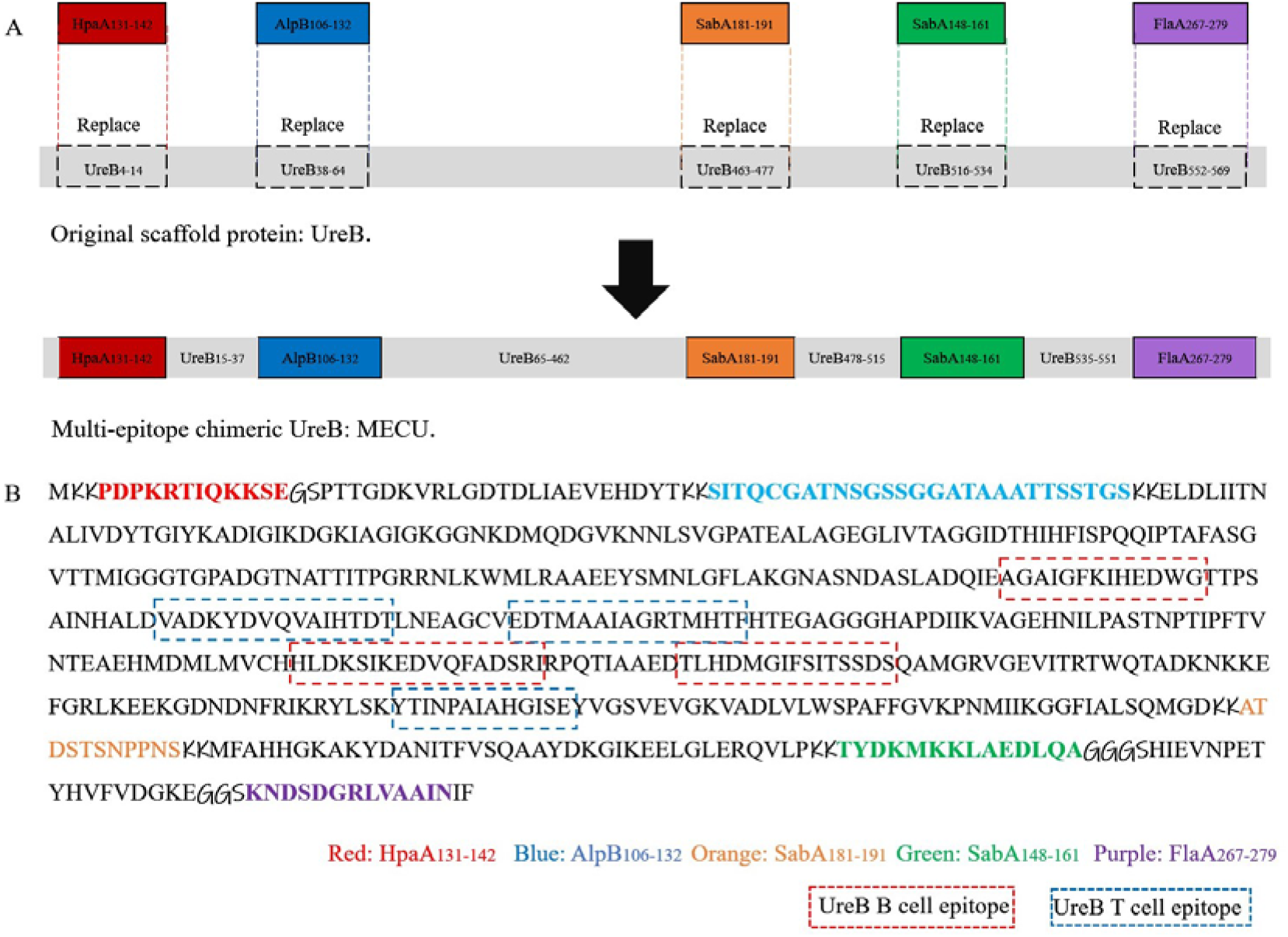
(A) Schematic representation of MECU construction. (B) The sequence of constructed MECU. Five peptide sequences in UreB were replaced by epitopes of HpaA, AlpB, SabA, and FlaA with linkers “KK”, “GS”, and “GGS”. The inherent B cell epitope sequences of UreB were marked by red dotted box and the inherent T cell epitope sequences of UreB were marked by blue dotted box. Besides, the epitope sequences from multiple antigens were marked by different colors.

Subsequently, the 3D structure of MECU was predicted by I-TASSER Server, a modeling structure prediction tool (Fig. 2A). In order to evaluate the structural reliability of the predicted structure, ProSA-web, RAMPAGE, and Verify 3D severs were used. The Z-score of MECU structure calculated by ProSA-web was −7.12, which is in the range of native protein conformation scores (Fig. 2B). Ramachandran plot from RAMPAGE showed that 92.5 % of residues were in favored region; 6.0 % of residues were in allowed region and 1.5 % of residues in outlier region (Fig. 2C). Verify 3D results showed that 86.83% of the residues have averaged 3D-1D score = 0.2 (Fig. 2D). All these results indicated that the predicted structure of MECU was reasonable and reliable.

**Fig. 2.**
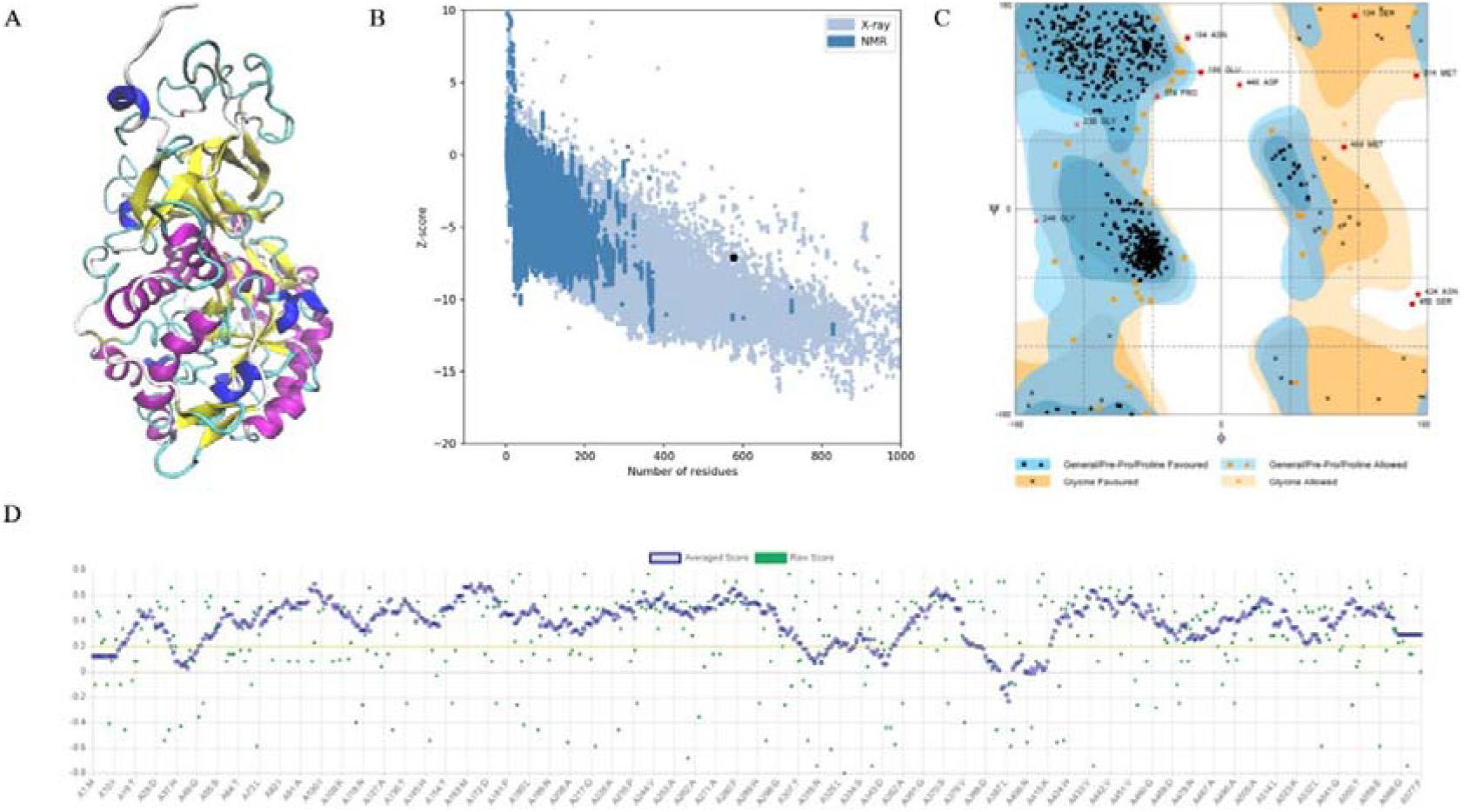
Tertiary structure prediction and validation of vaccine protein MECU. (A) Tertiary structure of MECU predicted by I-TASSER. (B) The z-score plot of the predicted structure by ProSA-web. Z-score= −7.12. (C) Ramachandran plot analysis of the predicted structure. Number of residues in favored region: 92.5 %; Number of residues in allowed region: 6.0 %; Number of residues in outlier region: 1.5%. (D) Verify3D analysis of the predicted structure. 86.83% of the residues have averaged 3D-1D score = 0.2.

Further, the structural alignment of MECU and UreB was performed by TM-align, which indicated that the structure of MECU was highly similar to that of UreB (RMSD = 0.62, Fig. 3A). The secondary structures of exogenous epitopes and replaced regions of UreB were analyzed for evaluating the design rationality. Both the B cell epitopes and replaced peptides contained the flexible loop fragment, whose substitutions at UreB-terminal had little effect on the stability of the whole skeleton (Fig. 3B). It could be seen that the chimeric epitopes were displayed on the surface of the MECU structure, which was beneficial to antibody binding (Fig. 3C).

**Fig. 3.**
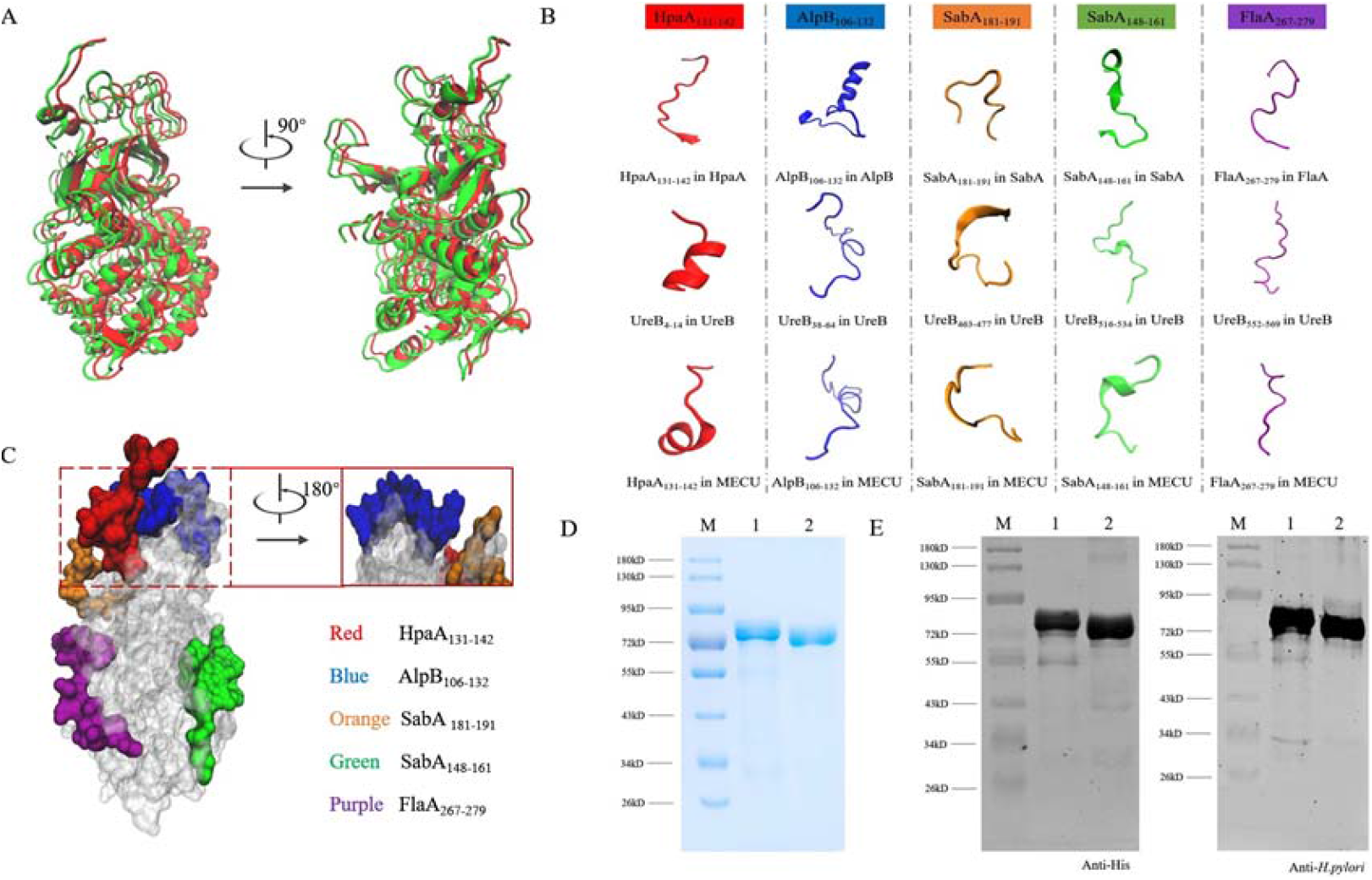
The design strategy and production of MECU. (A) Structural alignment of MECU (Red) and UreB (Green). RMSD=0.62. (B) The secondary structural changes of chimeric epitopes from the original antigens to MECU. (C) Epitope map of MECU. (D) Expression and purification of MECU visualized by SDS-PAGE. M: Maker; 1, UreB; 2, MECU.(E) Identity verification of MECU using Western blotting. M: Maker; 1, UreB; 2, MECU.

The MECU protein was successfully expressed and purified, whose molecular weight was similar to that of UreB (Fig. 3D). Both MECU and UreB protein could be recognized by mouse anti-His polyclonal antibody and rabbit anti-*H. pylori* polyclonal antibody (Fig. 3E).

### 3.2 Broadly reactive antibodies generated by MECU immunogen

To evaluate the actual effect of MECU design initially, mice were subcutaneously immunized by MECU for assay of antigen-specific antibodies in antisera. MECU could induce high levels of multiple antibodies specific to *H. pylori* antigens (UreB, FlaA, AlpB, SabA, and HpaA) while UreB could only induce high levels of antibodies to itself (Fig. 4A). It indicated that the epitopes from FlaA, AlpB, SabA, and HpaA have good immunogenicity and immunoreactivity, and MECU is a multivalent vaccine. In addition, the insertion of exogenous epitopes didn’t affect the humoral immunity of UreB in MECU due to the similar levels of antibodies specific to UreB.

**Fig. 4.**
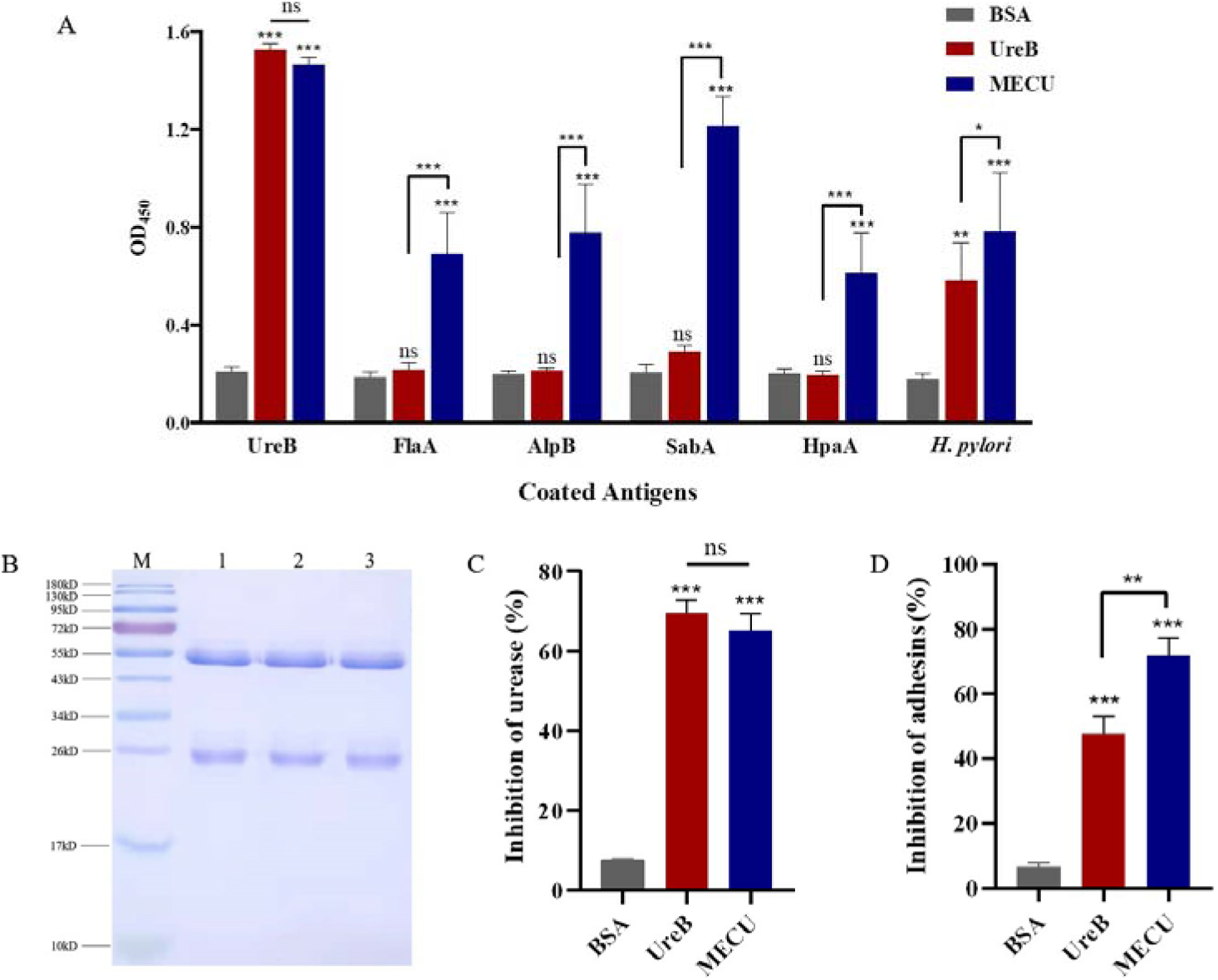
Evaluation of antibody response generated by MECU. (A) Determination of antigen-specific antibody in serum by ELISA after subcutaneous immunization with MECU, UreB, or BSA. (B) Visualization of IgG purified from serum of mice immunized by MECU, UreB, or BSA. M, Maker; 1, BSA; 2, UreB; 3, MECU (C) *H. pylori* urease neutralization test of the purified IgG. (D) Adherence inhibition assay of the purified IgG. These results were verified in triplicate assays. ***p < 0.001, **p < 0.01,*p < 0.05, ns, not significant.

To assay the neutralizing antibodies against *H. pylori* urease and adhesins, mouse IgG in antisera was purified and identified by SDS-PAGE (Fig. 4B). Anti-MECU IgG or anti-UreB IgG could inhibit the enzyme activity of *H. pylori* urease. However, the IgG induced by BSA had no significant inhibition (Fig. 4C). Besides, anti-MECU IgG or anti-UreB IgG could inhibit *H. pylori* adhesin to AGS cells, while anti-BSA IgG had no significant inhibition (Fig. 4D). Importantly, anti-MECU IgG had better adhesion inhibition to AGS cells than anti-UreB IgG.

### 3.3 Construction of CTB-MECU for oral immunization

According to the schematic representation (Fig. 5A), CTB-MECU immunogen was constructed by the addition of CTB to N-terminal of MECU with a flexible linker “DPRVPSS” and the construction of CTB-UreB was used as a reference. The CTB-MECU and CTB-UreB proteins were successfully expressed and purified (Fig. 5B). They could be identified by mouse anti-His polyclonal antibody and rabbit anti-*H. pylori* polyclonal antibody (Fig. 5C). Furthermore, the adjuvanticity of CTB in CTB-MECU was analyzed by GM1-ELISA. CTB-MECU and CTB-UreB were both able to bind the coating GM1, even though their binding abilities were weaker than the positive control CTB (Fig. 5D). UreB, MECU or the negative control BSA could not bind GM1 without the addition of CTB.

**Fig. 5.**
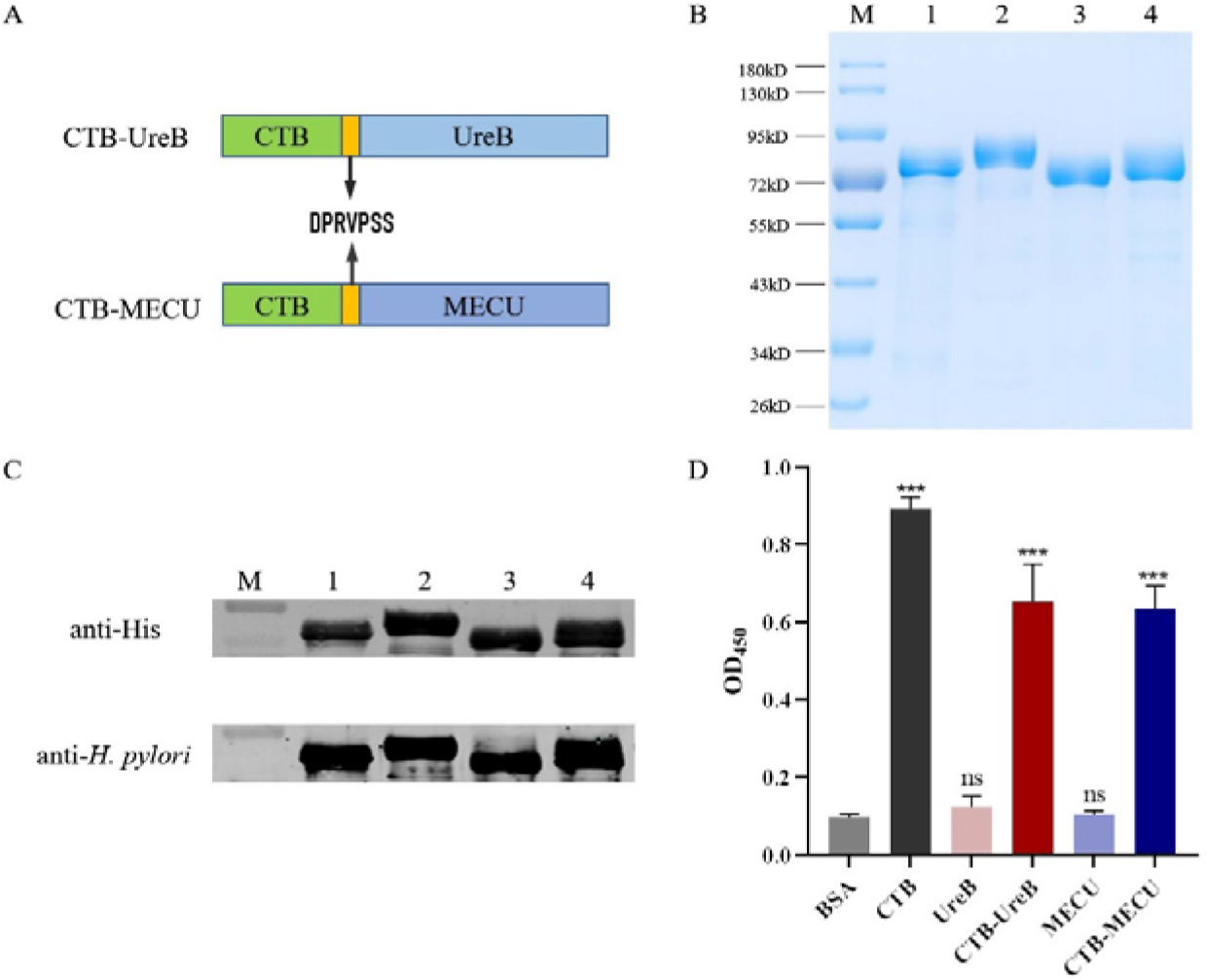
Construction of CTB-MECU for oral immunization. (A) Schematic representation of CTB-MECU construction with CTB-UreB as a reference. (B) Expression and purification of CTB-MECU visualized by SDS-PAGE. M, Maker; 1, UreB; 2, CTB-UreB; 3, MECU; 4, CTB-MECU. (C) Immunoreactivity of CTB-MECU probed by mouse anti-His polyclonal antibody and rabbit anti-*H. pylori* polyclonal antibody. M, Maker; 1, UreB; 2, CTB-UreB; 3, MECU; 4, CTB-MECU. (D) The adjuvant effect of CTB in CTB-MECU by GM1-ELISA. These results were verified in triplicate assays. ***p < 0.001, ns, not significant.

### 3.4 Evaluation of therapeutic vaccination

To evaluate the therapeutic effect of the constructed vaccine, MECU and UreB were vaccinated with FA intraperitoneally or with CTB intragastrically according to the procedure of therapeutic vaccination (Fig. 6A). The specific IgG and SIgA antibodies against *H. pylori* lysates were analyzed by ELISA after therapeutic vaccination. The measurement of *H. pylori* lysates-specific IgG antibodies in serum showed that mice immunized with MECU or UreB by intraperitoneal or oral route elicited significantly higher levels of IgG than the control (CTB+PBS+FA) group (Fig. 6B). In general, intraperitoneal vaccination induced higher levels of specific IgG antibodies than the oral route, which was significant in both UreB and MECU group. In addition, intraperitoneal vaccination with MECU induced higher levels of specific IgG antibodies than that with UreB, which was no difference in oral immunization. Oral immunization with MECU or UreB plus CTB remarkably increased the levels of SIgA antibodies against *H. pylori* lysates compared to the control group (Fig 6C). Moreover, CTB-MECU induced higher levels of SIgA antibodies against *H. pylori* lysates than CTB-UreB. Intraperitoneal vaccination with MECU or UreB did not induce significant SIgA antibodies against *H. pylori* lysates compared to the control group.

**Fig. 6.**
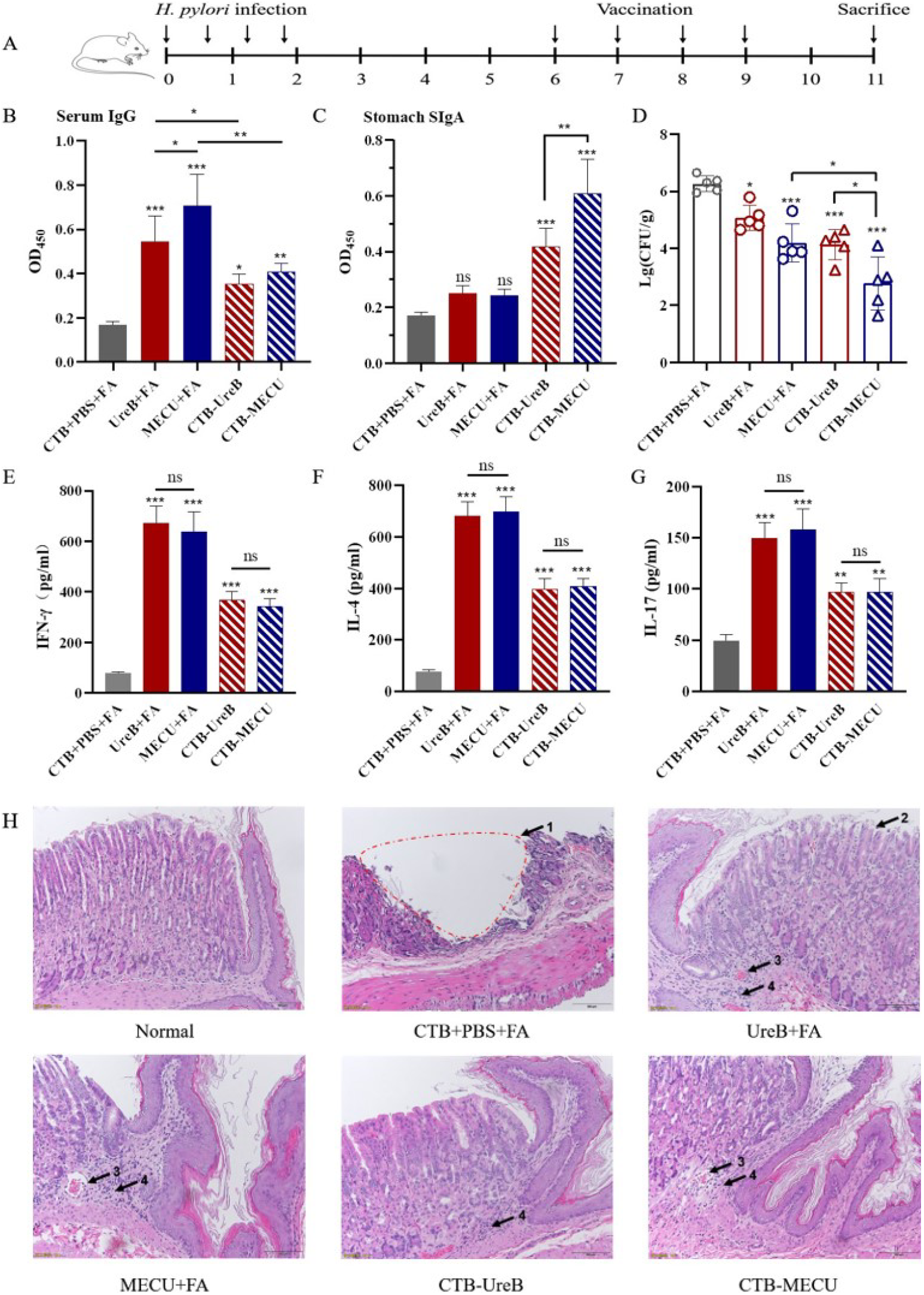
Evaluation of therapeutic vaccination. (A) The procedure of infection and therapeutic vaccination. Determination of serum IgG (B) or stomach SIgA (C) against *H. pylori* lysates after therapeutic vaccination (n=5). (D) Quantification of gastric *H. pylori* colonization by CFU counting. IFN-γ (E), IL-4 (F), and IL-17 (G) production in splenic lymphocytes stimulated by *H. pylori* lysates after therapeutic vaccination. (H) Gastric histology after oral therapeutic vaccination (HE stain). 1, Gastric ulcer; 2, Mucosal epithelial cells exfoliate and become necrotic, forming erosion; 3, Hemangiectasis; 4, Inflammatory cell infiltration. ***p < 0.001, **p < 0.01,*p < 0.05, ns, not significant.

The *H. pylori* colonization in the stomach was analyzed by quantitative culture. Compared with the control group, both two immunization routes with UreB or MECU significantly decreased the *H. pylori* loads in the stomachs (Fig 6D). Oral immunization had a better reduction of bacterial burden than intraperitoneal immunization, which was significant in MECU groups. Besides, MECU vaccination performed better at reducing bacterial colonization than UreB vaccination in oral immunization routes.

Further, the relevant cytokines IFN-γ, IL-4, and IL-17 in the supernatants of splenic lymphocyte cultures were determined after stimulation with *H. pylori* lysates using ELISA. *H. pylori* lysates significantly induced high levels of IFN-γ (Fig. 6E), IL-4 (Fig. 6F), and IL-17 (Fig. 6G) in splenic lymphocytes from mice immunized with UreB or MECU in both two immunization routes, but not those from the control group. There were no significant differences at the levels of IFN-γ, IL-4, and IL-17 cytokines between UreB and MECU group in each immunization route. However, the levels of IFN-γ, IL-4, and IL-17 in splenic lymphocytes from mice with intraperitoneal vaccination were higher than those with oral vaccination.

Therapeutic effect of MECU was also analyzed by histopathological analysis of stomach tissue. A severe stomach ulcer was found in the stomach from mice immunized with CTB and PBS plus FA. Erosive gastric epithelium was found in the stomach from mice immunized with UreB plus FA. Moderate or mild levels of inflammatory cell infiltration were found in the stomach immunized with UreB plus FA, MECU plus FA, CTB-UreB, or CTB-MECU (Fig. 6H).

## 4. Discussion

Urease, a key functional protein that helps *H. pylori* to colonize in the stomach, has become an important target for immunotherapy. The strongly immunogenic subunit UreB is a widely used antigen in the *H. pylori* vaccine studies^39^. However, the immune response induced by a single antigen is still limited. In this study, we constructed a multi-epitope chimeric antigen based on UreB structure, named MECU. Five positions on the surface of UreB structure were selected for displaying the exogenous B cell epitopes from FlaA, AlpB, SabA and HpaA. The replacement sites of exogenous epitopes did not affect inherent B or T cell epitopes of UreB, which means that UreB is not only a scaffold protein, but also retains its original strong immunogenicity (Fig. 1).

To further evaluate the design rationality of MECU, the constructed sequence was submitted to I-TASSER server for structural prediction. The structural alignment of UreB and MECU showed that the replacement of exogenous epitopes did not significantly change the principal skeleton of UreB structure (RMSD = 0.62). The exogenous epitopes could be displayed on the surface of MECU, which means that they have apparent accessibility to generate the humoral immune responses. Both the replaced peptides and the inserted B cell epitopes mostly contain flexible loop structures, which could ensure the stability of MECU structure to some extent.

After subcutaneous immunization, MECU induced a broader antibody response against antigens including FlaA, AlpB, SabA, and HpaA compared to UreB (Fig. 4A). In addition, the level of anti-*H. pylori* antibody in the antiserum from mice immunized with MECU was increased compared to that with UreB. Similar trends could be observed at the levels of anti-*H. pylori* IgG from mice with intraperitoneal immunization (Fig. 6B) and anti-*H. pylori* SIgA from mice with oral immunization (Fig. 6C). Besides, the purified anti-MECU antibodies could significantly inhibit the adhesion of *H. pylori* to AGS cells compared to anti-UreB antibodies (Fig. 4D). These results indicated that the exogenous B cell epitopes in MECU achieved the desired effect.

On the one hand, after immunogen engineering, MECU still induced a similar level of anti-UreB antibody to UreB (Fig. 4A). So did the anti-*H. pylori* urease neutralizing antibodies (Fig. 4C). On the other hand, there were no significant differences at the levels of IFN-γ, IL-4, and IL-17 cytokines between UreB and MECU group in both two immunization routes (Fig. 6E-G). This indicated that the inherent B and T cell epitopes of UreB were not affected by the replacements of exogenous epitopes. The original immunogenicity of UreB remained in MECU.

Both intraperitoneal vaccination and oral vaccination could significantly reduce the *H. pylori* colonization in the stomach and oral vaccination is more effective (Fig 6D). It results from that mucosal immunity could induce high levels of *H. pylori*-specific SIgA, the most abundant immunoglobulin of the mammalian mucosa (Fig 6C). In fact, the ability to induce significant levels of SIgA is a priority for the development of vaccine immunogens against gastrointestinal pathogens. The vaccines against *H. pylori* were mostly vaccinated intragastrically or nasally to induce mucosal immunity and produce high levels of specific SIgA^40^. In addition, CTB, a safe and efficient mucosal adjuvant, could enhance the levels of mucosal immunity^41^. However, intraperitoneal vaccination without inducing significant levels of *H. pylori*-specific SIgA could also reduce *H. pylori* colonization in the stomach, which was less effective than oral vaccination (Fig. 6C,D). It is possible to correlate the protection achieved with high levels of *H. pylori*-specific IgG (Fig. 6B). Some studies revealed that IgG in the murine intestine leads to the elimination of the Shiga toxin-producing *Escherichia coli*, virulent *Citrobacter rodentium*, and rotavirus^27, 42, 43^. The vaccines against *H. pylori* were also vaccinated by systemic immune routes such as intramuscular immunization^44, 45^. Consequently, our results and those reported by others support the idea that the effector functions of IgG in the defense against gastrointestinal pathogens shouldn’t be ignored.

In conclusion, we developed a promising formulation based on a recombinant chimeric protein named MECU, which was produced by the embedding of exogenous B cell epitopes on the surface of UreB structure. It could generate a broadly reactive antibody response and reduce *H. pylori* colonization in the murine stomach. Our study supports that the recombinant chimera containing epitopes of different antigens of *H. pylori* has the prospect of becoming an effective immunotherapy to control *H. pylori* infection.

